## Supplementary Information for "The C-terminus of the prototypical M2 muscarinic receptor localizes to the mitochondria and regulates cell respiration under stress conditions"

**This PDF file includes:**

Supplementary Text
Figs. S1 to S10
Tables S1 to S3
References (1 to 14)

### Supplementary results related to the muscarinic $M_2$ receptor

**Ligand binding properties of** $M_2$ **receptor mutants with single and double stop codons**

To understand the significance of the retained binding activity of the $M_2$stop228, verify whether the sequence downstream of the receptor is indeed translated and characterize the role played by the position of the stop codon in conferring this capacity to the receptor, we analyzed a number of receptor constructs with a stop codon placed either upstream or downstream of the i3 loop (**Fig. 1A** and **Fig. S1 and S3**).

None of the two $M_2$ receptor mutants bearing the stop codon in TM V and VI, such as $M_2$stop196 and $M_2$stop400, proves capable of binding [$^3$H]NMS (**Table S1** ). However, binding could still be rescued by co-transfecting $M_2$stop196 and $M_2$trunk(1-283) together. Similarly, binding of $M_2$stop400 could be restored by co-transfecting it with $M_2$tail(M-281-466) (**Table S1**). These observations are in line with previous data showing that receptor fragments interact with defective mutants [1]. Furthermore, they also corroborate the hypothesis that the carboxyl terminal both of $M_2$stop228 and $M_2$stop196 are indeed translated, resulting in an interaction with the truncated fragments.

To understand why mutants bearing a stop codon in regions TMV, TMVI and i3 loop could be rescued by co-transfection, and gain new insight into the mechanism underlying their interaction, new receptor mutants containing two additional stop codons, namely $M_2$stop196/stop400 and $M_2$stop228/stop400, were created (**Fig. S3**). When transfected alone in COS-7 cells (**Table S1**), none of the above receptor mutants proved capable of binding [$^3$H]NMS. Nevertheless, co-transfection of $M_2$stop228/stop400 and $M_2$tail(M-281-466) rescued the [$^3$H]NMS binding activity up to a value of Bmaxs comparable to that already observed by co-transfecting $M_2$trunk(1-283) with $M_2$tail(M-281-466) (**Table S1, and Table 1**). Conversely, co-transfection of $M_2$stop196/stop400 with any of the $M_2$ fragments, $M_2$trunk(1-283) and $M_2$tail(M-281-466), did not result in [$^3$H]NMS binding (**Table S1**). These data are indicative of the possibility that rescue of these receptor mutants depends entirely on the position of the stop codons. This is clearly demonstrated by the finding that the receptor cannot be rescued if the stop codons are present both in TMV and TMVI, as in $M_2$stop196/stop400. Based on these results, it also appears not possible that during transfection plasmid sequences could undergo homologous recombination[2] .

**Excluding stop codon read-through, termination-reinitiation and alternative splicing in the translation of $M_2$tail**

To rule out the possibility that the stop codon could be interpreted as a sense codon thus encoding an amino acid, a mechanism known as stop codon read-through[3, 4] , a frame shift was induced by inserting a sequence of four bases (AATT), fifteen nucleotides downstream of the stop codon 228: the mutant is hereafter named $M_2$stop228/fr.sh (**Fig. S3**). Our data show that the occurrence of a frame shift following insertion of this artificial stop codon does not alter [$^3$H]NMS binding activity of the mutant that essentially retains the Bmax values of 47 pmol/mg of protein (**Table S1**).

To check whether termination-reinitiation, a process known to occur in ribosomes translating the downstream Open Reading Frame (ORF) upon termination[5], was the mechanism responsible for the functional properties of the mutant $M_2$stop228, a 42 bases long palindromic sequence was inserted 15 nucleotides downstream of the stop codon (**Fig. S3**). When transcribed into mRNA, this palindromic sequence is expected to form a hairpin structure with a $\Delta G$ of −64 kcal/mol, a value sufficient to block ribosome scanning[6]. Our data show that insertion of this hairpin loop after the stop codon reduced slightly the [$^3$H]NMS binding activity of the receptor, attaining a Bmax value of 37 pmol/mg protein (**Table S1**). This evidence implies that termination-reinitiation is not the mechanism that accounts for [$^3$H]NMS binding of $M_2$stop228. These results were replicated by using similar mutants of the muscarinic $M_3$ receptor (**Table S2 and S3, see below**).

To verify whether a correct reading frame could be restored by the removal of the stop codon, through a mechanism of alternative splicing, mRNA was extracted from cells expressing $M_2$ and $M_2$stop228 and subjected to reverse transcriptase (see Materials and Methods). As shown in **Fig. S2B**, PCR amplification of the reverse transcript was resolved and detected with gel electrophoresis and showed a single band comparable in size to both $M_2$ and $M_2$stop228 receptor mRNAs, as it was for the control $M_2$ cDNA plasmid.

**Ligand binding properties of $M_2$stop228/stop248, $M_2$stop228/stop296, and $M_2$stop228/stop368**

To find out if any of the in-frame start codons was responsible for the translation of the carboxyl terminal fragment of the $M_2$stop228 receptor, the three ATG codons in frame after the stop codon 228 (**Fig. S1**), were each mutated to an additional stop codon. This yielded three new mutants, hereafter referred to as $M_2$stop228/stop248, $M_2$stop228/stop296, and $M_2$stop228/stop368 (**Fig. S3**). The first two mutants proved still capable of binding [$^3$H]NMS with $M_2$stop228/stop248 and $M_2$stop228/stop296, having Bmaxs of 49 fmol/mg and 38 fmol/mg of protein, respectively (**Table S1**). On the contrary, suppression of the third ATG in $M_2$stop228/stop368 abolished [$^3$H]NMS binding completely (**Table S1**), even though this capacity could be rescued by co-transfecting it together with the $M_2$tail(M-281-466) construct (**Table S1**).

**Preliminary analysis of the bicistronic Sirius-H-$M_2$i3(15n/30n)-EGFP construct to individuate the nucleotides required for internal ribosome entry**

The sequence alignment of the 33 nucleotides, from 1072 to 1104 of the $M_2$ i3 loop, with regions adjacent to the second ($M_1$), fourth ($M_3$ and $M_4$) and first ($M_5$) in frame ATGs of the other muscarinic receptors, revealed some conserved nucleotides, specifically an A in position 1078 and the three nucleotides AAG in position1090-1092 (four nucleotide AAGA 1090-1093 if we consider the two most related muscarinic $M_2$ and M4 receptors); furthermore the position of the ATG was conserved in four out of five muscarinic receptors (**Fig. S4C**).

Mutation of the conserved nucleotide A1078 with T, as in the Sirius-H-$M_2$i3(15n/30n)Mut#1-EGFP construct, reduced by half the EGFP fluorescence intensity compared to Sirius-H-$M_2$i3(15n/30n)-EGFP (**Fig. 2C**). Mutation of the four nucleotides AAGA1090-1093 with TTTT, as in the Sirius-H-$M_2$i3(15n/30n)Mut#2-EGFP construct, sharply decreased EGFP expression, retaining only a residual fluorescence of 24 ± 14% as respect to Sirius-H-$M_2$i3(15n/30n)-EGFP (**Fig. 2C**). Control experiments with mutants where non-conserved nucleotides were replaced, specifically Sirius-H-$M_2$i3(15n/30n)Mut#3-EGFP and Sirius-H-$M_2$i3(15n/30n)Mut#4-EGFP did not alter significantly EGFP expression (**Fig. 2C**).

Eventually we decided to alter the sequence that contains the residues that affected more dramatically EGFP expression. To this end, we mutated in T the sequence comprised between nucleotides 1083 and 1101, as in the Sirius-H-$M_2$i3(15n/30n)Mut#5-EGFP mutant. The nucleotide AAGA (1090-1093) was then reinserted in this mutant as in the Sirius-H-$M_2$i3(15n/30n)Mut#6-EGFP construct. As shown in **Fig. 2C**, only 12.3 ± 10.3% of the green fluorescence originally present in Sirius-H-$M_2$i3(15n/30n)-EGFP was retained in the Sirius-H-$M_2$i3(15n/30n)Mut#5-EGFP mutant. This loss, however, could be recovered up to 118 ± 7% of the Sirius-H-$M_2$i3(15n/30n)-EGFP level by re-inserting the four AAGA nucleotides into its sequence as in the Sirius-H-$M_2$i3(15n/30n)Mut#6-EGFP construct (**Fig. 2C**).

**Localization of the $M_2$-Muscarinic Receptor C-terminal upon cell stress**

We studied the expression of $M_2$tail(368-466)-mRuby2 – originated as a segment of the $M_2$-mRuby2 construct – and of $M_2$tail(368-466)-EGFP – originated as a segment of the $M_2$-i3-

tail-EGFP part – both derived from the $M_2$-mRuby2-STOP-$M_2$-i3-tail-EGFP mega construct, after serum starvation. **Fig. S9A-E** shows representative images of the mitochondrial localization of $M_2$tail(368-466)-EGFP and $M_2$tail(368-466)-mRuby2, in HEK293 cells stained with Mitotracker following two hours of starvation in PBS. As it can be clearly seen, expression of both $M_2$tail(368-466)-EGFP and $M_2$tail(368-466)-mRuby2 is visible in mitochondria, whereas at the plasma membrane only a $M_2$-mRuby2 signal is visible but none of the EGFP signal.

To further validate this observation, we repeated the exact same experiment where the mega construct was mutated in the third in frame methionine, resulting in $M_2$(M368A)-mRuby2-STOP-$M_2$-i3-tail-EGFP (**Fig. S3**). This time we did not observe expression of $M_2$tail(368-466)-mRuby2, and therefore no mRuby2 signal was visible in mitochondria **Fig. 9F-L**. On the other hand, $M_2$tail(368-466)-EGFP was still observed within mitochondria.

**Influence of the $M_2$-muscarinic receptor C-terminal fragment on mitochondrial oxygen consumption measured with the Clarke electrode**

The extent of oxygen consumption in COS-7 cells expressing the wild-type $M_2$, $M_2$stop228 and $M_2$tail(368-466) was also measured with the Clarke electrode. Our data show that no difference could be revealed, in term of total oxygen consumption, between cells transfected with the wild-type $M_2$ receptor or mock transfected cells (**Fig. S10B,C**). However, a sizable decrease of oxygen consumption rate was instead revealed, as compared to controls, in cells co-transfected with $M_2$stop228 and $M_2$tail(368-466) (**Fig. S10B,C**).

By comparison, cells transfected with the $M_2$trunk(1-228) fragment, did not change significantly their oxygen consumption rate. In order to evaluate whether $O_2$ depletion is coupled to oxidative phosphorylation, the ATP synthase inhibitor oligomycin, at the concentration of 17 nM, was added to the culture medium. The observation that oligomycin addition caused O2 consumption rate to decrease consistently in control samples, proved that the O2 consumption rate was coupled to oxidative phosphorylation (**Fig. S10B**), with an average value of 45 ± 6%. Oligomycin addition reduced also the O2 consumption rate of COS-7 cells transfected with $M_2$ and $M_2$trunk(1-228), while it only slightly reduced oxygen consumption rate in cells transfected with $M_2$stop228 and $M_2$tail(368-466) (**Fig. S10B,C**), thus suggesting that C-terminal-$M_2$ peptide suppresses only that portion of the O2 consumption rate that is coupled to the oxidative phosphorylation. It is worth nothing that in this assay the wild type $M_2$ receptor did not inhibit O2 consumption at variance with the Seahorse assay. The most parsimonious hypothesis to explain this discrepancy could be that in the assay with the Clarke electrode cells were not starved then the cap dependent translation is prevailing.

### Supplementary methods

#### Cloning and molecular biology

The two expression plasmids referred as $M_2$trunk(1-283) (containing transmembrane domains I-V and the N-terminal portion of the third cytoplasmic loop) and $M_2$tail(281-466) (containing transmembrane domains VI and VII, and the C-terminal portion of the third cytoplasmic loop) were described previously[7].

**Constructs for radioligand binding and immunoblotting**

$M_2$stop228 – This plasmid was created by substituting codon 228 (CAA) with a stop codon (TAA) at the N-terminal of the i3 loop of the wild type $M_2$ receptor.

$M_2$stop228/stop400 – This plasmid was created by inserting an additional stop codon – TGG was substituted with TAA – at the position codon 400 of the TM region VI of the $M_2$stop228 mutant.

$M_2$stop400 – This plasmid was created by substituting codon 400 (TGG) with a stop codon (TAA) in the TM region VI of the wild type $M_2$ receptor.

$M_2$stop196 – This plasmid was created by substituting codon 196 (TAT) with a stop codon (TAA) in the TM region V of the wild type $M_2$ receptor.

$M_2$stop196/stop400 – This plasmid was created by inserting an additional stop codon in the TM region VI of the $M_2$stop196 mutant, where TGG was substituted with TAA at position of the codon 400.

$M_2$stop228/fr.sh. – This plasmid was created by inserting four bases (AATT) fifteen nucleotides downstream of the stop codon to create a shift in the correct reading frame of the $M_2$stop228 mutant.

$M_2$stop228/hairpin – This construct was created by inserting a 42 bases long sequence (AGGGGCGCGTGGTGGCGGCTGCAGCCGCCACCACGCGCCCC), fifteen nucleotides downstream of the stop codon of the $M_2$stop228 mutant. Upon transcription, this palindromic sequence has been shown to form a hairpin structure with a ΔG value of −64 kcal/mol that blocks effectively ribosome scanning[6, 8].

$M_2$stop228/stop248, $M_2$stop228/stop296 and $M_2$stop228/stop368 – These three constructs were obtained by substituting the three in-frame ATG codons (1. ATG 248, 2. ATG 296 and 3. ATG 368) with the stop codon TAA downstream of the stop228 of the $M_2$stop228 mutant.

$M_2$trunk(1-228) – This plasmid was created by substituting codon 228 (CAA) with a stop codon (TAA) at the N-terminal of the i3 loop of the wild type $M_2$ receptor followed by removal of the downstream sequence.

$M_2$tail(368-466) – This construct was created by removing all bases of the $M_2$ receptor sequence up to codon 368 (ATG) of the C-terminal of the i3 loop. The resulting plasmid encodes for a polypeptide fragment that contains the trans-membrane domains VI and VII along with the C-terminal portion of the third cytoplasmic loop.

$M_2$(M368A) – In this construct the third in-frame methionine of the i3-loop of the $M_2$ muscarinic receptor (ATG) was mutated to alanine (GCG).

$M_2$-Myc, $M_2$stop228-Myc, $M_2$stopM368A-Myc, $M_2$tail(368-466)-Myc and $M_2$stop40-Myc – $M_2$-Myc was purchased from OriGene. This plasmid encodes the human $M_2$ receptor with a Myc (tag at the C-terminus of the protein. $M_2$stop228-Myc was obtained by replacing the 0.9 Kb BmtI-PspOMI fragment of the $M_2$stop228 mutant with the corresponding fragment of $M_2$-Myc. $M_2$stop400-Myc was obtained by substituting the codon 400 (TGG) with a stop codon (TAA) of the $M_2$-Myc. All the constructs have an additional DDK tag after the C-Myc. Subsequently, all C-Myc constructs were extracted by PCR and subcloned into a bicistrionic pVitro2-MCS plasmid (Invivogen), allowing for the expression of a reporter gene (in our case the red fluorescent protein mRuby2) after an IRES sequence, in order to check for transfection efficiency.

**Generation of bicistronic constructs expressing Sirius and EGFP proteins** (**Fig. 2A**)

Sirius[PacI-$M_2$i3(685-1101)]EGFP – This bicistronic construct was created by inserting the i3 loop of the muscarinic $M_2$ receptor, from nucleotide 685 to nucleotide 1101, between the ultramarine fluorescent protein (Sirius) and the green fluorescent protein (EGFP). Eight nucleotides corresponding to the recognition site of the PacI enzyme were inserted between Sirius and the i3 loop of the wild type $M_2$ receptor to alter the reading frame downstream of the Sirius stop codon. Hereafter this plasmid is referred to as Sirius-$M_2$i3(417n)-EGFP.

Sirius[PacI-$M_2$i3(685-699)-Hairpin-G]EGFP – This bicistronic construct was created by inserting a 42 nucleotides hairpin loop (see above) between Sirius and the EGFP fluorescent protein. The hairpin loop was spaced from the Sirius stop codon by inserting a PacI recognition sequence and 15 nucleotides of the $M_2$ i3 loop sequence comprised between nucleotide 685 and nucleotide 699. A G nucleotide was also inserted following the 42 nucleotides of the hairpin loop and upstream of the initial ATG triplet of the EGFP to restore a correct reading frame. Throughout the text we referred to this plasmid as Sirius-H-EGFP.

Sirius[PacI-$M_2$i3(685-699)-Hairpin-G-PacI-$M_2$i3(685-1101)]EGFP – This bicistronic construct was created by adding 417 nucleotides of the i3 loop of $M_2$ from nucleotide 685 to nucleotide 1101 to the plasmid Sirius[PacI-$M_2$i3(685-699)-

Hairpin-G]EGFP. An additional PacI restriction site was also inserted upstream of this segment of the i3 loop. Throughout the text we referred to this plasmid as Sirius-H-$M_2$i3(417n)-EGFP.

Sirius[PacI-$M_2$i3(685-699)-Hairpin-G-PacI-$M_2$i3(685-699/1072-1101)]EGFP – This bicistronic construct was created by deleting from the plasmid Sirius[PacI-$M_2$i3(685-699)-Hirpin-G-PacI-$M_2$i3(685-1101)]EGFP 372 nucleotides of the $M_2$ i3 loop, from nucleotide 700 to nucleotide 1071. Throughout the text we referred to this plasmid as Sirius-H-$M_2$i3(15n/30n)-EGFP.

Sirius[PacI-$M_2$i3(685-699)-Hirpin-G-PacI-$M_2$i3(685-699)]EGFP – This bicistronic construct was created by adding 15 nucleotides of the i3 loop of $M_2$ from nucleotide 685 to nucleotide 699 to the plasmid Sirius[PacI-$M_2$i3(685-699)-Hirpin-G]EGFP. An additional PacI restriction site was also inserted upstream of this short segment of the i3 loop. Throughout the text we will refer to this plasmid as Sirius-H-$M_2$i3(15n)-EGFP.

Sirius[PacI-$M_2$i3(685-699)-Hirpin-G-PacI-$M_2$i3(685-1071)]EGFP – This bicistronic construct was created by adding 387 nucleotides of the i3 loop of $M_2$ form nucleotide 685 to nucleotide 1071 to the plasmid Sirius[PacI-$M_2$i3(685-699)-Hirpin-G]EGFP. An additional PacI restriction site was also inserted upstream this long segment of the i3 loop. Throughout the text we will refer to this plasmid as Sirius-H-$M_2$i3(387n)-EGFP.

Mutants of the Sirius-H-$M_2$i3(15n/30n)-EGFP plasmid – Based on the sequence alignment of amine and adenosine GPCRs, several conserved nucleotides were mutated in order to define the putative IRES sequence. These mutants are reported in **Fig. 2A**.

The fluorescence intensities of the two spectral bands were quantified by the use of a fluorometer and the EGFP intensity values normalized to those of Sirius.

**Constructs for fluorescence microscopy**

$M_2$-EGFP – The wild type $M_2$ muscarinic receptor was cloned into the pEGFP-N1 expression cassette, between the restriction sites HindIII and XbaI.

$M_2$-mRuby2 – The wild type $M_2$ muscarinic receptor was cloned into the mRuby2 vector (Addgene plasmid #40260).

$M_2$tail(368-466)-EGFP – The $M_2$tail(368-466)-EGFP was constructed so that the receptor protein would start from the third in-frame methionine of the $M_2$ i3 loop (M368). The EGFP gene was fused C-terminally following a short restriction site sequence (TCTAGA) of the XbaI enzyme.

$M_2$tail(368-466)-mRuby2 – The $M_2$tail(368-466)-mRuby2 was constructed so that the receptor protein would start from the third in-frame methionine of the $M_2$ i3 loop (M368). The mRuby2 gene was fused C-terminally following a short restriction site sequence (TCTAGA) of the XbaI enzyme.

$M_2$-mRuby2-STOP-$M_2$-i3-tail-EGFP – This mega construct was obtained by fusing two preceding constructs, $M_2$-mRuby2 and $M_2$tail(368-466)-EGFP, in a unique plasmid, but the $M_2$tail started at codon 229 ($M_2$tail(229-466)-EGFP). After the stop codon of mRuby2 and before the codon 229 of $M_2$tail was inserted a short restriction site sequence (TCCGGA) of the BspEI enzyme (**Fig. 3D**).

$M_2$(M368A)-mRuby2-STOP-$M_2$-i3-tail-EGFP – This mega construct is analogous to $M_2$-mRuby2-STOP-$M_2$-i3-tail-EGFP, but the third in-frame methionine of the i3-loop of the $M_2$ muscarinic receptor was mutated to alanine (M368A) (**Fig. S3**).

$M_2$fr.sh-EGFP – A single base insertion (G) upstream of nucleotide 1102 of the wild type $M_2$ receptor, just before the third in-frame methionine of the i3-loop (M368), induces a frame shift that, following 2 aminoacids, generates a stop codon

(TAA) in the amino acid position 370 of the new reading frame. The construct is then fused to EGFP.

$M_2$fr.sh-mRuby2 – As above, but the EGFP is replaced by a mRuby2

$M_2$-hairpin-EGFP – The 3$^{rd}$ i3 loop of the $M_2$ receptor (between resides 228 and 338) was replaced by the sequence AGGGGCGCGTGGTGGCGGCTGCAGCCGCCACCACGCGCCCCt, generating a mRNA hairpin, as discussed above before the 30 nt sequence upstream of M368.

$M_2$-GFP11 – A short peptide 16 aminoacids long from the GFP protein, GFP11, was fused to the C-terminus of the muscarinic $M_2$ receptor. GFP11 was custom syntethised and inserted as a linker between the XbaI and XhoI restriction sites in the plasmid backbone of $M_2$-EGFP.

Mito-GFP1-10 – The mitochondrial targeting sequence of the cytochrome c oxidase subunit 8A (COX8A) was fused to the N-terminal of GFP1-10 fragment of GFP. GFP1-10 was purchased from Addgene as Addgene plasmid 70219.

Mito-GFP11 – The mitochondrial targeting sequence of COX8A was fused to GFP11 fragment. Mito sequence was obtained from Addgene plasmid 23348.

SMAC-GFP1-10 – The mitochondrial targeting sequence of the SMAC protein was fused to the N-terminal of GFP1-10 fragment. SMAC was obtained from Addgene plasmid 40881.

SMAC-GFP11 – The mitochondrial targeting sequence of the SMAC protein was fused to GFP11 fragment.

$M_2$tail(368-466)-GFP1-10 – The GFP1-10 fragment was fused to the C-terminus of the muscarinic $M_2$tail(368-466) receptor fragment.

$M_2$tail(368-466)-GFP11 – The GFP11 fragment was fused to the C-terminus of the muscarinic $M_2$tail(368-466) receptor fragment.

SMAC-mCitrine – mCitrine was fused to the C-terminus of the mitochondrial targeting sequence of SMAC using restriction enzymes: BamHI and NotI. SMAC sequence was obtained from Addgene plasmid 40881. mCitrine originates from plasmid FLAGmMOR-mCitirine, in pcDNA3.1.

Mito-mCitrine - mCitrine was fused to the C-terminus of the mitochondrial targeting sequence of COX8A using restriction enzymes: BamHI and NotI. Cox8A (Mito) sequence was obtained from Addgene plasmid 23348.

CV-mCitrine - mCitrine was fused to the C-terminus of the mitochondrial targeting sequence of Complex V (CV) using restriction enzymes: EcoRI and NotI. CV sequence was obtained from Addgene plasmid 213884.

$M_2$tail-mTurquoise2 - mTurquoise2 was fused to the C-terminus of $M_2$tail using the restriction enzymes: XbaI and NotI. mTuquoise2 originates from the plasmid pc-FLAG-mTq2-b2AR, in pcDNA3.1.

**Constructs for in-vitro mitochondrial import assays**

PSP64-$M_2$tail(368-466)-5Met - The vector backbone PSP64 (courtesy of Ruaridh Edwards). The following primers: FWD: ATGATGTAAtctagaggaggcggacgc, REV: CATCATCATccttgtagcgcctatgttcttataatg

were used to add five methionines to the C-terminal domain of the $M_2$tail(368-466)-GFP11 template (see above) using Q5® Site-Directed Mutagenesis Kit (New England Biolabs® Inc.). The construct was then extracted with appropriate restriction enzymes (HindIII and XbaI) and ligated into the vector backbone PSP64 (courtesy of Ruaridh Edwards).

PSP64-$M_2$R-5Met – The following primers: FWD: ATGATGTAAtctagaggaggcggacgc REV: CATCATCATccttgtagcgcctatgttcttataatg were used to add five methionines to the C-terminal domain of to the $M_2$-GFP11 template (see above) using Q5® Site-Directed Mutagenesis Kit (New England Biolabs® Inc.). The construct was then extracted with appropriate restriction enzymes (HindIII and XbaI) and ligated into the vector backbone PSP64.

All constructs were sequenced either with a Genetic Analyzer 3500 (Applied Biosystem®) or by a professional sequencing service (LGC Genomics, Berlin).

**Immunoblot of muscarinic $M_2$ receptor mutants transfected in HEK293 cells**

Cells were seeded in 10 cm plates and transfected after 24h according to manufacturer protocols (Effectene, Qiagen) using 2 µg of plasmid DNA. 48h after transfection cells were washed (2x) in ice cold PBS. 200 µL per plate of ice-cold lysis buffer were added. Lysis buffer was a RIPA buffer: 50 mM Tris-HCL pH 8, 150 mM NaCl, 1% TRITON, 0.5% sodium deoxycholate and 0.1% SDS. The following compounds were added to the buffer to a final 1x dilution: 100x Halt Protease Inhibitor Cocktail (Thermo Fisher), 100x 0.5 M EDTA and (10x) 10 mM PMSF. Cells were then scraped and the cell lysate transferred to pre-cooled 1.5 mL Eppendorf tubes. The tubes were placed in a Thermomixer at 4 °C, 300 rpm for 30 minutes. Then the Eppendorf tubes were centrifuged for 30 minutes at 14000 rpm at 4 °C. The supernatant was then transferred to new Eppendorf tubes. Protein concentration in the cell lysates was quantified by a BCA assay (Pierce BCA Protein Assay Kit, from Thermo Fisher) according to manufacturer instructions.

Cell lysates were loaded into 10% polyacrylamide gels. Cell lysates were loaded using Laemmli Buffer 2x (Sigma Aldrich) in equal mass. As a reference marker we loaded 5 uL of PageRuler Prestained Protein Ladder (Thermo Fisher). Gels were mounted into a Mini Protean Tetra Cell kit (BioRad) using 1x SDS running buffer. Gels were ran using a constant voltage of 90V for approximately 15 minutes, and afterwards at 130V for 45 minutes. Gels were transferred to PVDF membranes. Wet transfer was achieved by using the Mini Trans Blot Module (BioRad) in Wet Transfer Buffer (25 mM Tris pH 8.3, 192 mM Glycine and 20% MetOH) at 350 mA for 90 minutes. The immunoblotted membranes were then blocked in 5% Milk in TBS-T for 1 hour at room temperature. The membrane was then incubated overnight at 4 °C with primary antibody solution using the following concentrations depending on the antibody used: 1:1000 anti β-actin (13E5) rabbit primary antibody (Cell Signaling), 1:1000 anti myc-Tag (9B11) mouse primary antibody (Cell Signaling). After overnight incubation, the membranes were washed 3x 10 minutes in TBS-T and incubated at RT for 1 h with a secondary antibody (1:10000 anti-mouse IgG/(anti rabbit) HRP-linked antibody from Cell Signaling).

After the incubation with the secondary antibody, the membrane was washed 3x 10 minutes in TBS-T. Detection was achieved by eliciting HRP luminescence by incubating the membrane with SuperSignal WestFemto solution (Thermo Fisher) according to the manufacturer's instructions, and imaging the membrane using a c600 Transilluminator from Azure Biosystems.

**Immunoblot of muscarinic $M_2$ receptor transfected in COS-7 cells**

SDS-Triton protein extraction - COS-7 cells were collected 48 h after transfection and treated with lysis buffer (50 mM TrisHCl pH 7.8, 1% Triton X100, 0.1% SDS, 250 mM NaCl, 5 mM EDTA, 100 mM NaF, 2 mM NaPPi, 2 mM Na3VO4, 1 mM PMSF). Cell lysates were then centrifuged at 16000 g for 15 minutes at 4 °C and supernatants, containing solubilized receptors resolved by SDS-PAGE or stored at -80°C. Samples were mixed with 65 mM Tris, 10% glycerol, 2% SDS, 0.1 M DTT, 0.001% bromophenol blue, pH 6.80 with HCl, boiled for 5 min and applied to a 10% SDS-PAGE.

SDS-PAGE - Resolved proteins were transferred to a PVDF membrane (Bio-Rad). The membrane was blocked in blotto (5% non-fat dry milk in 1xTBS plus 0.1% Tween 20) and the expression level of the $M_2$ receptor was assayed with a mouse anti-Myc-tag diluted 1:1000 in blotto and thereafter in a HRP-conjugated secondary antibody diluted 1:1000 in blotto. Membranes were then incubated in SuperSignal West Pico chemiluminescent substrate (Thermo Fisher Scientific Inc.) and the bands detected using a Bio-Rad ChemiDoc XRSplus imaging system. Optical densities of blot bands were finally determined using a computer-assisted densitometer (ImageJ U. S. National Institutes of Health, Bethesda, Maryland, USA), normalized versus the tubulin internal control.

**Detection of phosphorylated ERK1**

On day zero, HeLa cells were plated on 6-well plates (70,000 cells/well). On day 1, the cells were transiently transfected with the plasmid of interest (1 µg DNA/well). On day 3, the cells were exposed to serum-free medium until the day of the assay. On day 4, the cells were treated with 100 μM carbachol (time 0'; 1'; 5' and 20') at 37 °C. The cells were then lysed in a buffer containing 50 mM TrisHCl (pH 7.8), 1% Triton X100, 0.1% SDS, 250 mM NaCl, 5 mM EDTA, 100 mM NaF, 2 mM NaPPi, 2 mM Na3VO4, 1 mM PMSF. Samples were incubated on ice for 30 min and then centrifuged at 17,000 rpm for 15 min at 4 °C. The supernatants were recovered and assayed for protein concentration. Protein extracts were run on a 10% SDS-PAGE and transferred on a PVDF membrane (Bio-Rad). The membrane was then blocked in blotto (5% non-fat dry milk in 1xTBS plus 0.1% Tween 20) and the extent of phosphorylation of (ERK) mitogen-activated protein kinase was determined by immunoblotting with anti-phospho-ERK (Sigma-Aldrich) diluted 1:1000 in blotto. The blots were stripped and re-blotted with the anti-ERK (Sigma-Aldrich), diluted 1:1000 in blotto, to estimate the total amount of kinase loaded. Detection of the immunoreactive bands was carried out by the enhanced chemiluminescence method (SuperSignal west Pico, Thermo Fisher Scientific Inc.), by using a ChemiDoc XRSplus imaging system (Bio-Rad Laboratories, Milan, Italy), and their optical densities determined by using a computer-assisted densitometer.

**Cell culture of H9c2 cells**

H9c2 (ATCC, H9c2 (ATCC; CRL-1446)) were cultured in DMEM (Dulbecco's Modified Eagle Medium (DMEM, 4.5 g/L D-glucose, 110mg/L Sodium Pyruvate) (from Thermofisher Scientific), supplemented with 10% Fetal Bovine Serum (FBS) and 1% Penicillin/Streptomycin (P/S). Cells were grown in 25 $cm^2$ (check) flasks and maintained at 37C and 5% $CO_2$. Cells were seeded into dimension here glass coverslips which had been coated using Poly-L-Lysine solution (PLL, 0.01%) in 6-well plates.

Cells were transfected using Lipofectamine 2000 (Thermofisher) reagent according to the manufacturer's protocol, such that 2.5 μg plasmid DNA was transfected per coverslip at least 24 hours before imaging. Cells were stained with MitoTracker Deep Red FM (Thermofisher) through a 100x dilution of 1mM MitoTracker to each coverslip. They were stained with Hoechst 33342 Solution through a 1000x dilution of 1mg/ml Hoechst to each coverslip.

**Oxygen consumption measured with the Clark electrode**

A Clark type electrode-based polarographic method was used to evaluate oxygen consumption, in a 2 ml volume chamber kept under continuous stirring at the constant temperature of 37 °C[9] . Briefly, after determination of the baseline of oxygen consumption in the presence of the sole culture medium, 6 x 106 of transfected cells were added, and the disappearance of oxygen was monitored through a Yellow Springs Instruments Model 53 Oxygen Monitor device (YSI Inc., Yellow Springs, OH). Since the electrode consumes oxygen during measurement, the rate of oxygen decrease in 2 ml of DMEM media with no cells was subtracted from the sample's oxygen consumption rates. Results are expressed as ng-atom of oxygen/min/106cells. For the evaluation of oxygen consumption rate in the absence of oxidative phosphorylation, the ATP synthase inhibitor oligomycin was added to the measurement vessel through a fine needle.

**$G_i$ protein activation FRET assays**

HEK/TSA cells were co-transfected with plasmids carrying the $M_2$ receptor or its mutants (Table), together with a plasmid carrying the $Gi_1$ FRET biosensor at a 1:3 ratio[10]. The co-transfection was performed using Effectene Transfection Reagent (Qiagen) following the manufacturers protocol.

| **Plasmid** | **$M_2$ receptor** | **Fluorescent reporter** |
|---|---|---|
| pVitro2_$M_2$wt_myc_IRES_mRuby2 | $M_2$wt | mRuby2 |

| | | |
|---|---|---|
| pVitro2_M2M368A_myc_IRES_mRuby2 | $M_2$M368A | mRuby2 |
| pVitro2_M2stop228_myc_IRES_mRuby2 | $M_2$stop228 | mRuby2 |

HEK/TSA cells were seeded in a 100mm dish to a total number of 2x106 cells in 10ml DMEM (Thermo Fisher Scientific) supplemented with 10% FCS, 1% glutamine and 1% penicillin/streptomycin (PenStrep). After 24 hours, the co-transfection was performed as described above. After additional 24 hours, transfection efficiency was checked using a fluorescence microscope. After confirmation of successful transfection, the cells were plated in a 96 well black bottom plate to a total number of 5x104cells per 200µl per well. Untransfected HEK/TSA were plated as a control. The next day the cells were washed three times with 200µl 1X Hanks' Balanced Salt Solution (HBSS, Thermo Fisher Scientific) and the FRET baseline was measured using the BioTek SynergyTM Neo2 Hybrid Multi-Mode Microplate Reader. The donor fluorophore was excited with a wavelength of 430/30 and the emission was measured at 491/30. The acceptor fluorophore was excited with a wavelength of 500/18 and the emission measured at 541/20. The FRET signal was measured with an excitation wavelength of 430/20 and an emission wavelength of 541/20.

Following the baseline measurement, 200µl acetylcholine diluted in 1X HBSS were added to the cells. The concentrations were chosen to be around the published EC50 of acetylcholine for the $M_2$ R of around 0,1-0,03µM. 200µl 1X HBSS were added to wells containing the untransfected control cells.

### Supplementary results related to the muscarinic $M_3$ receptor

#### Ligand binding and functional properties of co-transfected $M_3$trunk(1-272)/$M_3$tail(M-388-589) fragments

As previously shown by[7], none of the two muscarinic receptor fragments, generated by splitting the receptor at the level of the i3 loop, $M_3$trunk(1-272) and $M_3$tail(M-388-589) showed [$^3$H]NMS binding activity when expressed alone in COS-7 cells. In contrast, a considerable number of specific [$^3$H]NMS binding sites was observed after co-expression of $M_3$trunk with $M_3$tail, and the binding affinities for [$^3$H]NMS and carbachol were identical to the wild type $M_3$ receptors. Furthermore, co-expression of $M_3$trunk with $M_3$tail resulted in functional receptors able to increase phosphatidylinositol hydrolysis (**Table S2**).

**Ligand binding and functional properties of M3stop273**

In line with what as been observed with, $M_2$stop228, $M_3$stop273, a receptor mutant bearing a stop codon at the beginning of the i3 loop, exhibited binding activity when expressed in COS-7 cells, albeit at very low level of expression, such as 52.3 fmol/mg of protein. Remarkably, the calculated [$^3$H]NMS KD value, obtained from direct saturation experiments was similar to that calculated for the wild type $M_3$ receptor (**Table S2**). Furthermore, competition experiments showed that the agonist carbachol was able to inhibit [$^3$H]NMS binding and the isotherm was best fitted by a one site binding model (**Table S2**).

$M_3$stop273 was functionally active increasing the phosphatidylinositol hydrolysis after carbachol stimulation, even though the extent of response was reduced compared to the wild type $M_3$ receptor (**Table S2**).

**Lack of [$^3$H]NMS binding of $M_3$stop240 and $M_3$stop503**

In analogy to what has been done with the $M_2$ receptor, we analyzed $M_3$ receptor constructs with a stop codon upstream and downstream the i3 loop. None of the $M_3$ receptor mutants bearing the stop codon in TM V, such as M3stop240, or in TM VI, such as $M_3$stop503, were able to bind [$^3$H]NMS (**Table S3**). Nevertheless, the binding of $M_3$stop240 could be rescued by co-transfection with $M_3$trunk(1-272), while the binding of $M_3$stop503 was restored in the presence of $M_3$tail(M-388-589) (**Table S3**). These data agree with previous ones showing the capability of receptor fragments to interact functionally with defective mutants.

**Lack of [$^3$H]NMS binding of the double mutants $M_3$stop240/stop503 and $M_3$stop273/stop503**

In order to deepen our knowledge about the mechanism involved to rescue mutants bearing the stop codon in the regions TMV, (TMVI) and i3 loop, we created two additional mutants with two stop codons, such as $M_3$stop240/stop503 and $M_3$stop273/stop503.

None of these two receptors exhibited binding to [$^3$H]NMS when they were transfected alone in COS-7 cells (**Table S3**). Nevertheless, co-transfection of $M_3$stop273/stop503 with $M_3$tail(M-388-589) rescued [$^3$H]NMS binding activity, with Bmaxs similar to those observed with the co-transfection of $M_3$trunk(1-272) with $M_3$tail(M-388-589) (**Table S3**). Conversely, the co-transfection of $M_3$stop240/stop503 with any of the $M_3$ fragments did result in [$^3$H]NMS binding (**Table S3**).

**Ligand binding properties of $M_3$stop273/fr.sh**

As a stop codon could hypothetically be interpreted as a sense codon encoding for an amino acid, in order to check for stop codon read-through, a frame shift was created by inserting a four base AATT 15 nucleotides after the stop codon 273, resulting in the mutant named $M_3$stop273/fr.sh. . The presence of the frame-shift after the artificial stop codon did not alter the [$^3$H]NMS binding activity of the receptor that showed Bmax values of 51.5 pmol/mg of protein. (**Table S3**).

**Ligand binding properties of $M_3$stop273/hairpin**

In order to check if termination re-initiation was the mechanism responsible of the functional properties of the mutant $M_3$stop273, a 42 bases long palindromic structure, was inserted 15 nucleotides after the stop codon, resulting in the $M_3$stop273/hairpin receptor. This palindromic sequence when transcribed into mRNA, forms a hairpin structure with a ΔG value of −64 kcal/mol, which can effectively block ribosome scanning[8] . The presence of the hairpin loop after the stop codon, which avoids the re-initiation process, slightly reduced but did not abolish [$^3$H]NMS binding activity of the receptor, with Bmax values that was 29.7 pmol/mg of protein (**Table S3**).

### Supplementary methods related to the muscarinic $M_3$ receptor

#### Generation of mutant muscarinic $M_3$ receptors

Rat $M_3$ muscarinic receptor expressed in pcD plasmid[11] [11, 12][11, 12]was used to construct the different mutants of muscarinic $M_3$ receptors. The two expression plasmids referred as $M_3$trunk(1-272), containing transmembrane domains I-V and the N-terminal portion of the third cytoplasmic loop, and $M_3$tail(M-388-589), containing transmembrane domains VI and VII, and the C-terminal portion of the third cytoplasmic loop, were described previously [7].

$M_3$stop273 – This plasmid was created substituting codon 273 (CAA) at the N-terminal of the i3 loop of $M_3$ with a stop codon 273 (TAA).

$M_3$stop273/stop503 – This plasmid was created by inserting in $M_3$stop273 an additional stop codon in TM region VI, at position codon 503, TGG was substituted with TAA.

$M_3$stop503 – This plasmid was created by substituting codon 503 (TGG) in TM region VI of $M_3$ with a stop codon (TAA).

$M_3$stop240 – This plasmid was created by substituting codon 240 (TAC) in TM region V of $M_3$ with a stop codon (TAA).

$M_3$stop240/stop503 – This plasmid was created by inserting in $M_3$stop240 an additional stop codon in TM region VI, at position codon 503, TGG was substituted with TAA.

$M_3$stop273/fr.sh. – This plasmid was created by inserting in $M_3$stop273 a four bases AATT fifteen nucleotides after the stop codon in order to create a shift in the correct reading frame.

$M_3$stop273/hairpin – This construct was created by inserting in $M_3$stop273, fifteen nucleotides after the stop codon, a 42 bases long sequence (AGGGGCGCGTGGTGGCGGCTGCAGCCGCCACCACGCGCCCCT). This palindromic sequence when transcribed into mRNA, forms a hairpin structure with a $\Delta G$ value of −64 kcal/mol, which can effectively block ribosome scanning [6].

**Generation of a bicistronic construct expressing the i3 loop of muscarinic M3 receptor between the Sirius and EGFP proteins**

Sirius-$M_3$i3(558n)-EGFP – This plasmid was generated by inserting between the two fluorescent protein Sirius and EGFP the i3 loop of the muscarinic $M_3$ receptor, from nucleotide 880 to nucleotide 1437. A PacI enzyme was inserted between the stop codon of Sirius and the beginning of the $M_3$ i3 loop (**Fig. S5A**).

### Supplementary Figures and Tables

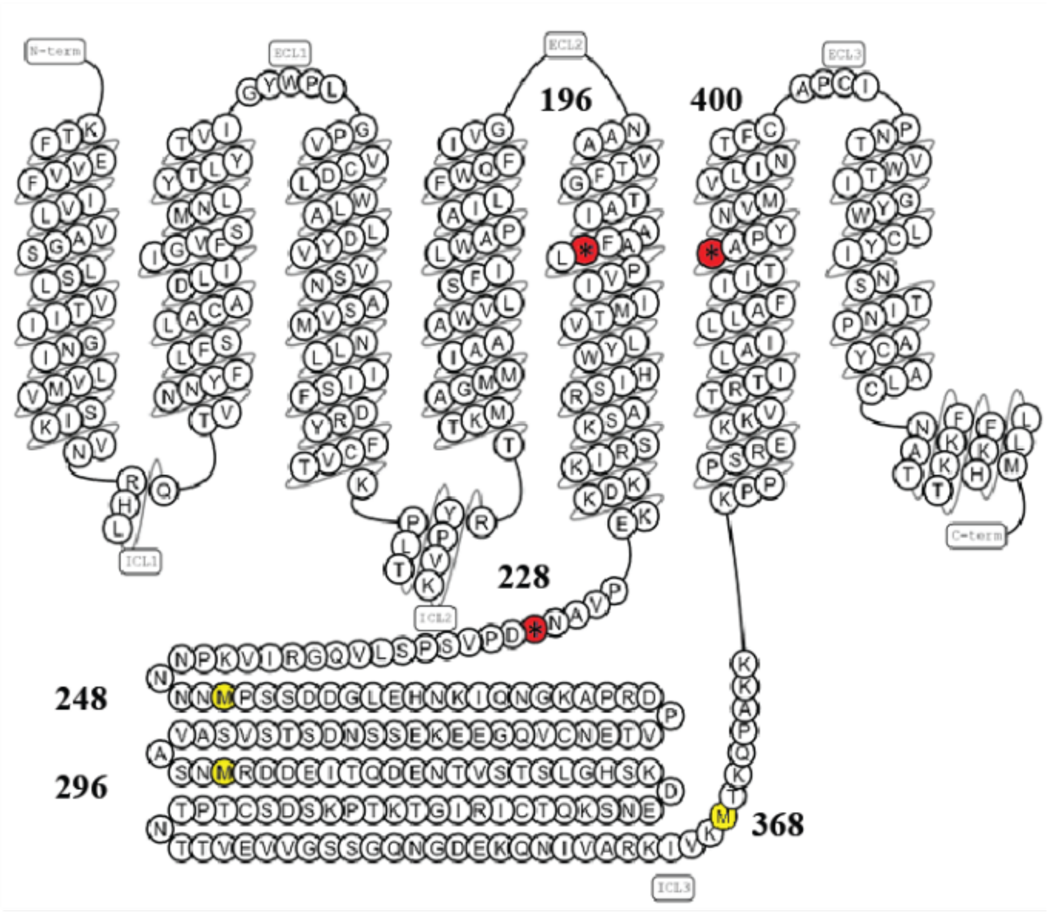


**Fig. S1. Snake diagram of the $M_2$ receptor.**

Key residues are highlighted[13]. Asterisks in the red circles denote the insertion of stop codons within transmembrane regions V (196) and VI (400), as well as the i3 loop (228). The in-frame methionine highlighted in yellow are the ones within the third loop that have been substituted with stop codons to investigate the initiation start site of $M_2$ Cterminal fragment.

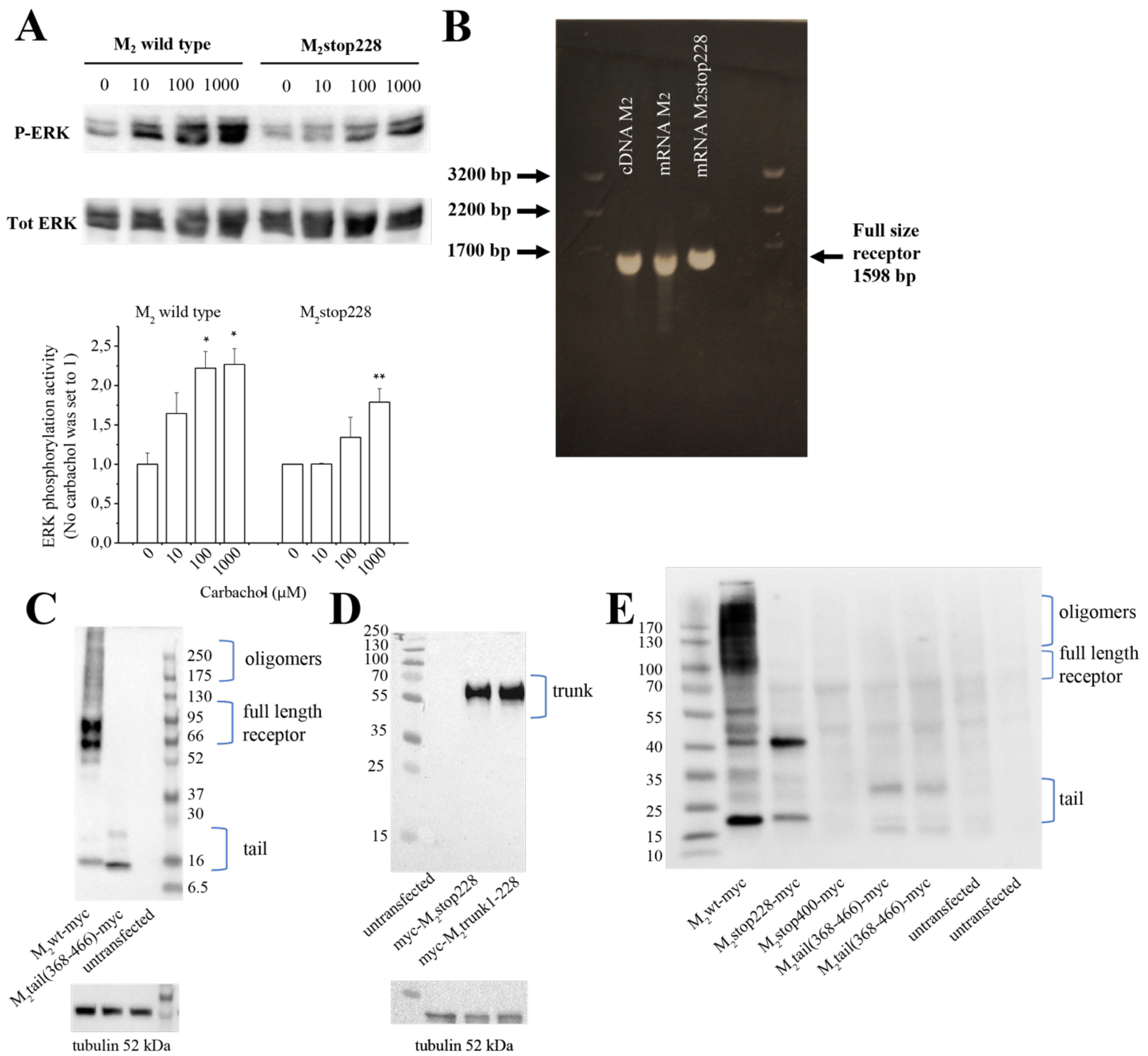


**Fig. S2. M2 receptor mutants: signalling, alternative splicing and Western blotting**.

**A,** Dose response curve of carbachol induced ERK phosphorylation in HeLa cells transiently transfected with $M_2$ and $M_2$stop228 muscarinic receptors. Cells were stimulated for 5 minutes with the indicated concentration of carbachol. Phosphorylated ERK (P-ERK) was normalized vs total ERK (Tot ERK). Significance values reported in the graphs were determined by a one-tailed Student t-test, p-values: ** $0.001<p<0.01$; * $0.01<p<0.05$ was calculated against carbachol 0 µM. **B,** mRNAs were extracted by COS-7 cells transfected with $M_2$ wild type and $M_2$stop228, subjected to reverse transcriptase and the resulting cDNA amplified by PCR with two oligos directed to the 5' and 3' end of the receptors. The gel shows, in both $M_2$ wild type and $M_2$stop228, a single band of 1401 bp corresponding to the full-size receptor, running at the same level of the band amplified directly from the $M_2$ pcD plasmid. **C,** Western blot of whole cell lysates of HEK293 cells. $M_2$-Myc and $M_2$tail(368-466)-Myc were transiently transfected in HEK293 cells and immunodetected via Western blot together with an untransfected control. Loading control was verified by immunoblotting tubulin. **D,** Western blot of whole-cell lysates untransfected HEK293 (lane 1) and then HEK293 cells transfected with myc-$M_2$Stop228 (Lane 2) and myc-$M_2$Trunk(1-228) (Lane 3). Loading control was verified by detecting tubulin. **E,** Western blot of whole cell lysates of COS-7 cells. $M_2$-Myc, $M_2$stop228-Myc, $M_2$tail(368-466)-Myc, $M_2$stop400-Myc were transiently transfected in COS-7 cells and immunodetected via Western blot together with an untransfected control. Source data for panel A can be found in S1 Data.

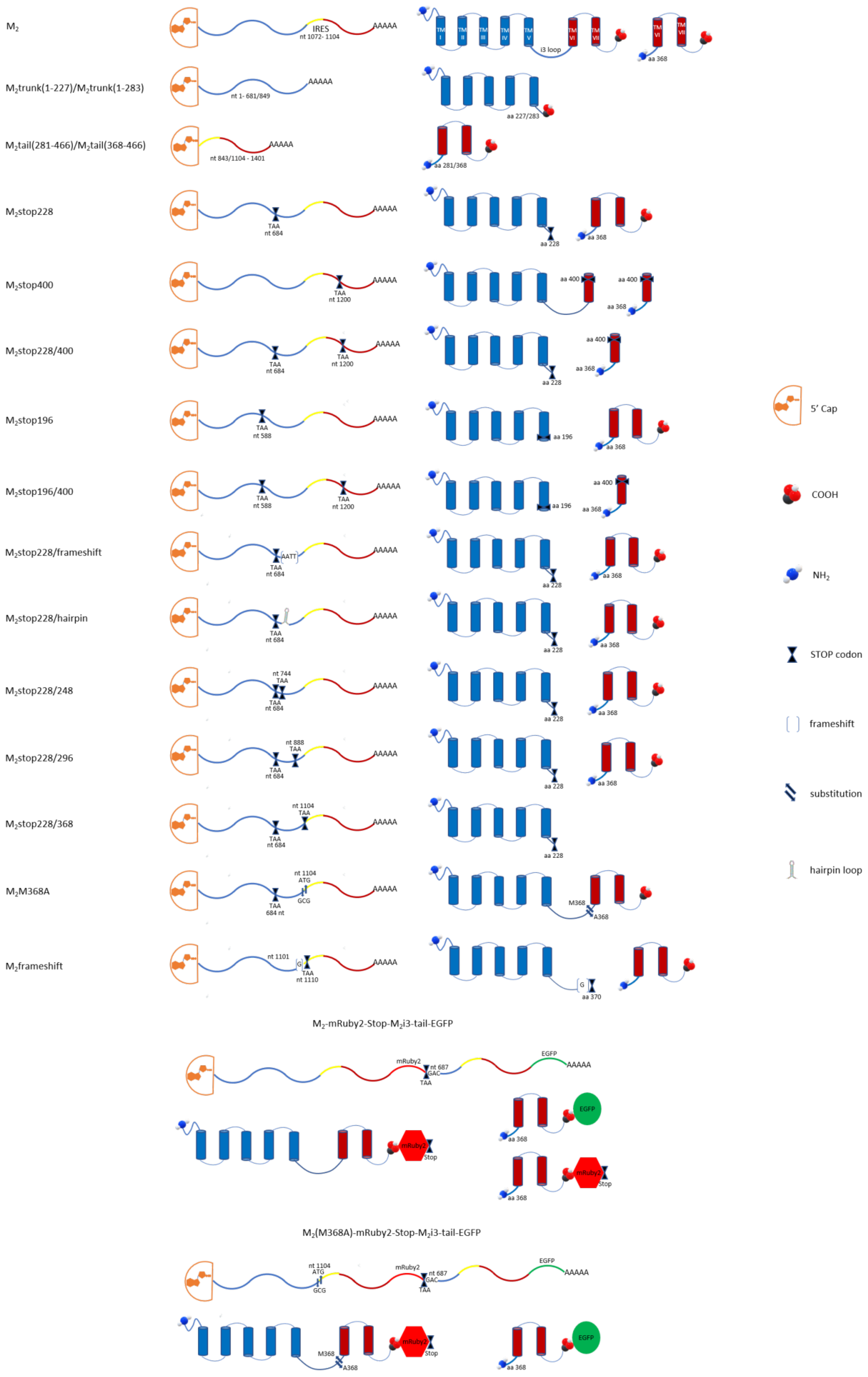
$M_2$
$M_2$trunk(1-227)/$M_2$trunk(1-283)
$M_2$tail(281-466)/$M_2$tail(368-466)
$M_2$stop228
$M_2$stop400
$M_2$stop228/400
$M_2$stop196
$M_2$stop196/400
$M_2$stop228/frameshift
$M_2$stop228/hairpin
$M_2$stop228/248
$M_2$stop228/296
$M_2$stop228/368
$M_2$M368A
$M_2$frameshift
$M_2$-mRuby2-Stop-$M_2$i3-tail-EGFP
$M_2$(M368A)-mRuby2-Stop-$M_2$i3-tail-EGFP
IRES
nt 1072- 1104
AAAAA
i3 loop
aa 368
nt 1- 681/849
aa 227/283
nt 843/1104 - 1401
aa 281/368
TAA
nt 684
aa 228
nt 1200
aa 400
nt 588
aa 196
AATT
nt 744
nt 888
nt 1104
ATG
GCG
684 nt
M368
A368
nt 1101
nt 1110
aa 370
mRuby2
nt 687
GAC
EGFP
Stop
5' Cap
COOH
$NH_2$
STOP codon
frameshift
substitution
hairpin loop

**Fig. S3. Schematic representation of wild type muscarinic $M_2$ (human) receptors and derived mutants.**

Each construct was obtained as described under **Materials and Methods**. The left column indicates the mRNA product from each construct, the right column the expected protein product. For the last two constructs, above mRNA product, and below the expected protein product.

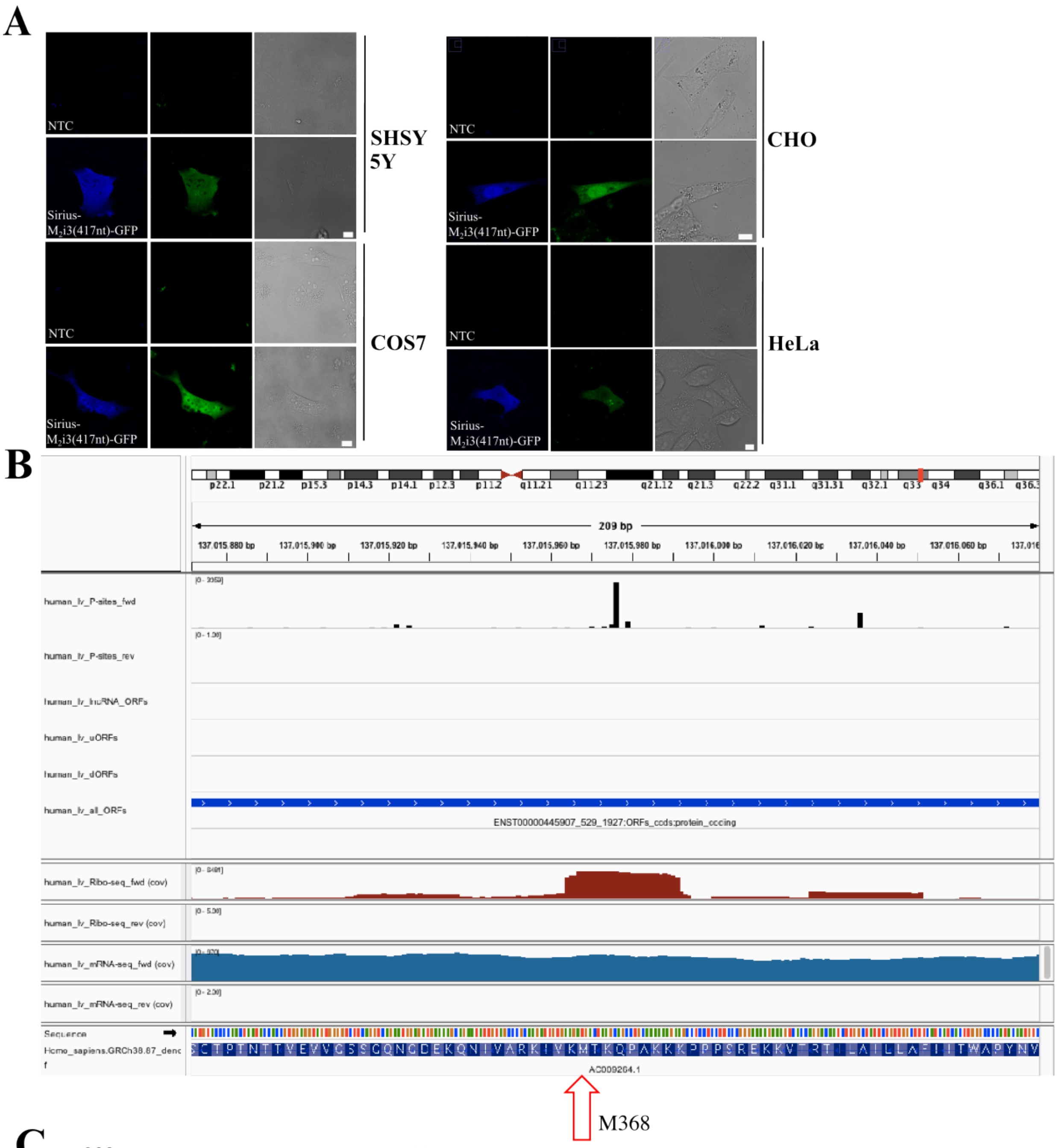


C

```
           992                                  1024
            ↓                                    ↓
CHRM2      CAGAATATTGTAGCCCGCAAGATTGTGAAGATG           ATG (Met368 in the i3 loop) is 57 nucleotides from TMD6
CHRM4      GCTCGCAACCAGGTGCGCAAGAAGCGGCAGATG           ATG (Met391 in the i3 loop) is 30 nucleotides from TMD6
CHRM1      GAGAGGAGCCTGGCTCCGAAGTGGTGATCAAGATG         ATG (Met304 in the i3 loop) is 186 nucleotides from TMD6
CHRM3      ACCAGAAGTCAGATCACTAAGCGGAAAAGGATG           ATG (Met481 in the i3 loop) is 30 nucleotides from TMD6
CHRM5      ACCAACAATGGCTGTCACAAGGTGAAAATCATG           ATG (Met403 in the i3 loop) is 120 nucleotides from TMD6
```

**Fig. S4. Expression of the bicistronic plasmid Sirius-M2i3(417n)-EGFP in four different cell lines, ribosome profiling data and sequence alignment**

**A,** Expression of Sirius was driven by the canonical scanning mechanism of translation initiation, while EGFP expression was driven by an IRES dependent mechanism. All Sirius-M2i3(417n)-EGFP transfected cells expressed both the Sirius and EGFP proteins. On average, 25 ± 4% of the COS-7 cells transfected with Sirius-$M_2$i3(417n)-EGFP plasmid were blue fluorescence positive, while 17 ± 2% were green fluorescent positive, with only a few cells being only green. Excitation conducted at 405 nm (410-450 nm detection) for Sirius, and at 488 nm (500-550 nm detection) for GFP. Scale bars are 10 µm throughout. **B,** Ribosome profiling data from heart-specific transcriptomic databases[14], where the $M_2$ receptor is highly expressed. When looking at left ventricle data, the ribosome coverage data is prominent in correspondence of the third i3loop in frame-methionine M368 (highlighted by the red arrow). In this case, also p-site hits are observed. **C,** Alignment of the nucleotide sequence 1072-1104 of the $M_2$ i3 loop with analogous sequences of the other four muscarinic receptors. The criteria for the alignment are described in the supplementary data. In yellow are highlighted the conserved nucleotides. In bold are the in frame ATG codons. Next to each sequence is indicated the codon number of the in-frame ATG and its distance in nucleotides from the beginning of TMDVI.

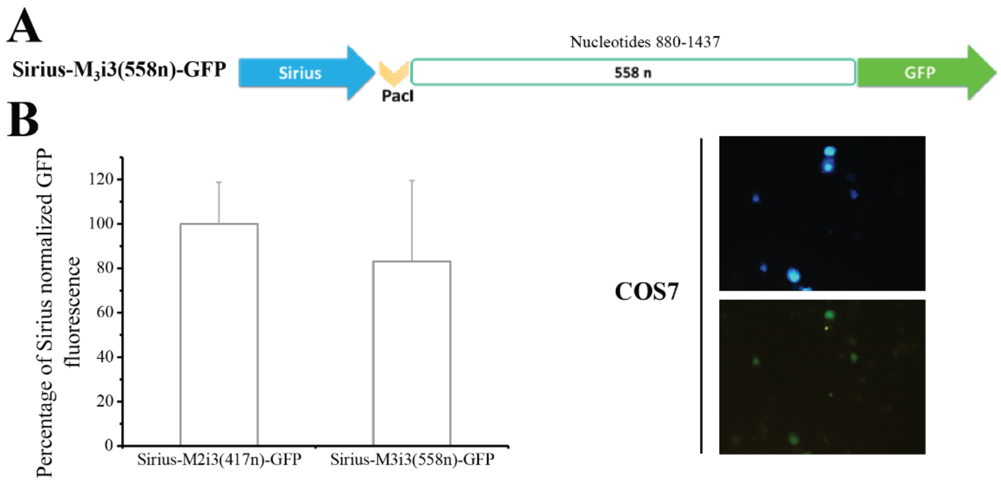


**Fig. S5. Expression of the bicistronic plasmid Sirius-M3i3(558n)-EGFP.**

**A,** Schematic representation of the bicistronic plasmid bearing the i3 loop of the rat muscarinic $M_3$ receptor (558 nucleotides from nucleotide 880 to nucleotide 1437) between the coding regions of the Sirius and EGFP fluorescent proteins. **B,** EGFP expression in COS-7 cells transfected with the bicistronic plasmid Sirius-$M_3$i3(558n)-EGFP. Source data for panel B can be found in S1 Data.

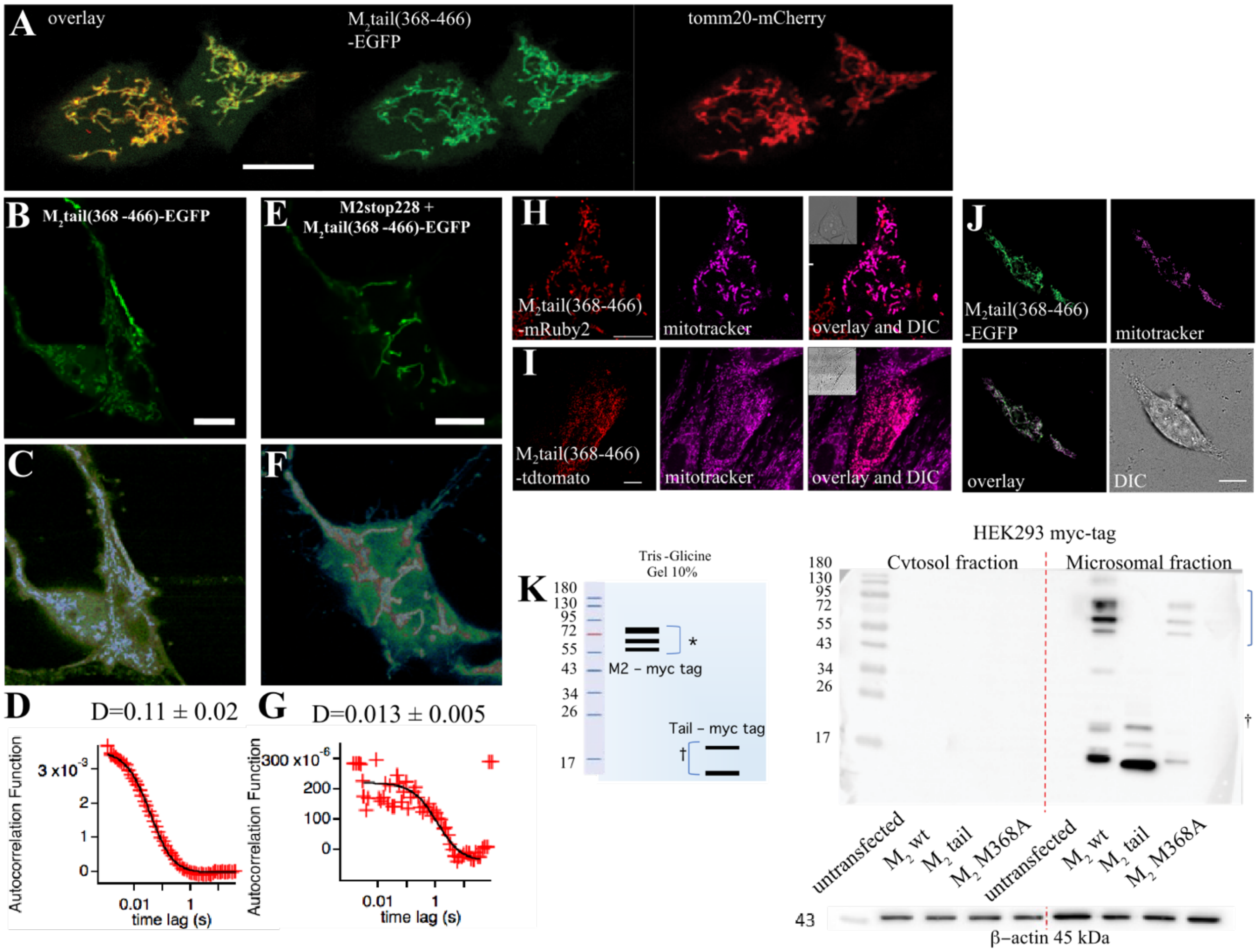


**Fig. S6. Subcellular localization of the $M_2$tail(368-466)**

**A,** Confocal image displaying the colocalization (left of $M_2$tail(368-466)-EGFP (green, middle) in HEK293 cells, co-expressed with the outer mitochondrial membrane marker Tomm20-mCherry (red, right). **B,** expression of $M_2$tail-EGFP displaying marked mitochondria and cytosolic localization. **C,** Glasbey colorscale representation of panel (B) highlighting the low-intensity pixels. **D,** autocorrelation function of the $M_2$tail(368-466)-EGFP diffusion taken at or in proximity of the basal membrane, yielding a mean diffusion coefficient of 0.11 μ$M_2$/s, incompatible with membrane diffusion. **E,** co-expression of $M_2$stop228 and $M_2$tail(368-466)-EGFP. $M_2$stop228 is unlabeled. $M_2$tail(368-466)-EGFP maintains the localization to the mitochondria observed in cells expressing $M_2$tail(368-466)-EGFP alone, but it is also localized to the plasma membrane, visible in panel **F,** Glasbey colorscale representation of panel E highlighting the low-intensity pixels. **G,** autocorrelation function of the $M_2$tail(368-466)-EGFP diffusion taken at or in proximity of the basal membrane, yielding a mean diffusion coefficient of 0.013 μ$M_2$/s, compatible with membrane diffusion. **H,** Cellular localization of $M_2$tail-mRuby2, together with Mitotracker deep red staining of the mitochondrial network and corresponding DIC image. **I,** Cellular localization of $M_2$tail-tdtomato and corresponding DIC image. **J,** Confocal sequential images displaying the localization of fluorescently labeled $M_2$tail(368-466)-EGFP (green), together with the mitochondrial network (magenta), colocalization (white) and DIC image in COS-7 cells. All imaging panels originate from Laser Scanning Confocal Microscope sequential acquisitions, with laser lines 488 nm (EGFP) 561 nm (mRuby2, mCherry and tdtomato) and 633 nm (Mitotracker deep red), and corresponding emission filters in the ranges 520-600 nm, 570-620 nm and 640-750 nm. Scale bars 10 µm. **K**, Western blot (10% TRIS-Glycine PAA gel) of the cytosolic and microsomal fractions resulting from the mitochondria purification of lysates of HEK293 transfected with myc-tagged constructs, as displayed in **Fig. 3C**. Loading control involved immunoblotting for $\beta$-actin. Source data for panels D and G can be found in S1 Data.

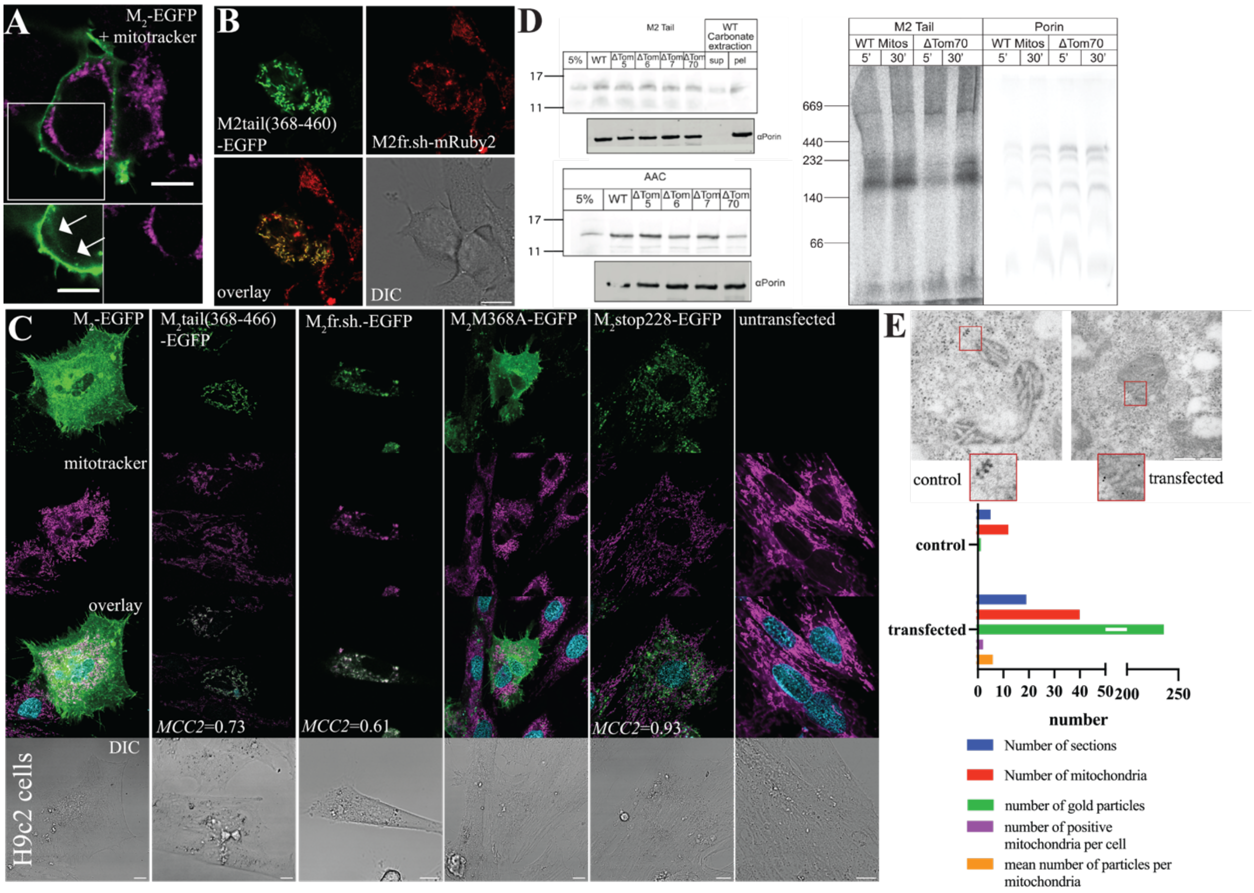


**Fig. S7. Mitochondrial localization of the M2tail fragment under endogenous IRES production and in vitro mitochondrial import of M2tail(368-466)**

**A,** Confocal image of $M_2$-EGFP (green) expressed in conjunction with Mitotracker (magenta) in HEK293 cells. Separate panels below. **B,** Confocal image of HEK293 cell co-expressing $M_2$tail(368-460) and $M_2$fr.sh-mRuby2, together with the merged fluorescence image and DIC. **C,** Confocal micrographs of representative H9c2 cells expressing $M_2$wt-EGFP, $M_2$tail(368-466)-EGFP, $M_2$fr.sh.M368A-EGFP, $M_2$M368A-EGFP, $M_2$stop228-EGFP and an untransfected control. Panels display, from top to bottom EGFP (green), Mitotracker (Magenta), overlay (EGFP, Mitotracker and, where present, Hoechst 33342 (Cyan)) and DIC (grays). Scale bars are 10 µm. Confocal sequential acquisitions were performed with 405 nm excitation and

420-460 nm detection (Hoechst), 488 nm excitation and 520-600 nm detection (EGFP), and 633 nm excitation and 650-750 nm detection (Mitotracker) using HyD detectors in Photon Counting Mode. Manders Correlation Coefficient $M_2$ (fraction of green/$M_2$tail features within magenta/mitochondria features) is indicated on the overlay images. **D,** 35S-labelled $M_2$tail(368-466) or AAC (an integral internal mitochondrial membrane protein) were imported into isolated yeast mitochondria originating from different knock-out lines for outer membrane transporters (ΔTom). ΔTom5=68% of WT, ΔTom6=87% of WT, ΔTom7=87% of WT, ΔTom70=59% of WT . Samples were incubated for 30 minutes, and then washed in breaking buffer. After import, 100 μg of mitochondria were subject to Carbonate extraction to determine if $M_2$tail(368-466) is integrated into a lipid bilayer (Pellet, 68%) or loosely associated/soluble (Supernatant, 32%). Samples were then loaded onto a 12% Tris-Tricine gel followed by semi-dry transfer and visualized via a phosphorimager (n=1). To test for equal mitochondrial loading anti-$\alpha$Porin was used via western blot. (right)35S-labelled $M_2$tail(368-466) and $\alpha$Porin were imported into isolated wild-type or ΔTom70 mitochondria for the indicated timepoints. Samples were then washed in breaking buffer and loaded on blue-native page followed by gel drying and visualized via a phosphorimager (n=1). **E,** Representative IEM micrographs of a control (left) and transfected (right) COS-7 cells. Scale bar is 1 μm. Statistics of IEM staining, both in transfected as well as control cells. The discernible 'dark spots' observed in the control, likely attributed to varied exposure settings during imaging, exhibit a noticeably larger size and less distinct shapes compared to the transfected samples, where circular, sharp, and intense gold nanoparticles are evident. Source data for panel E can be found in S1 Data.

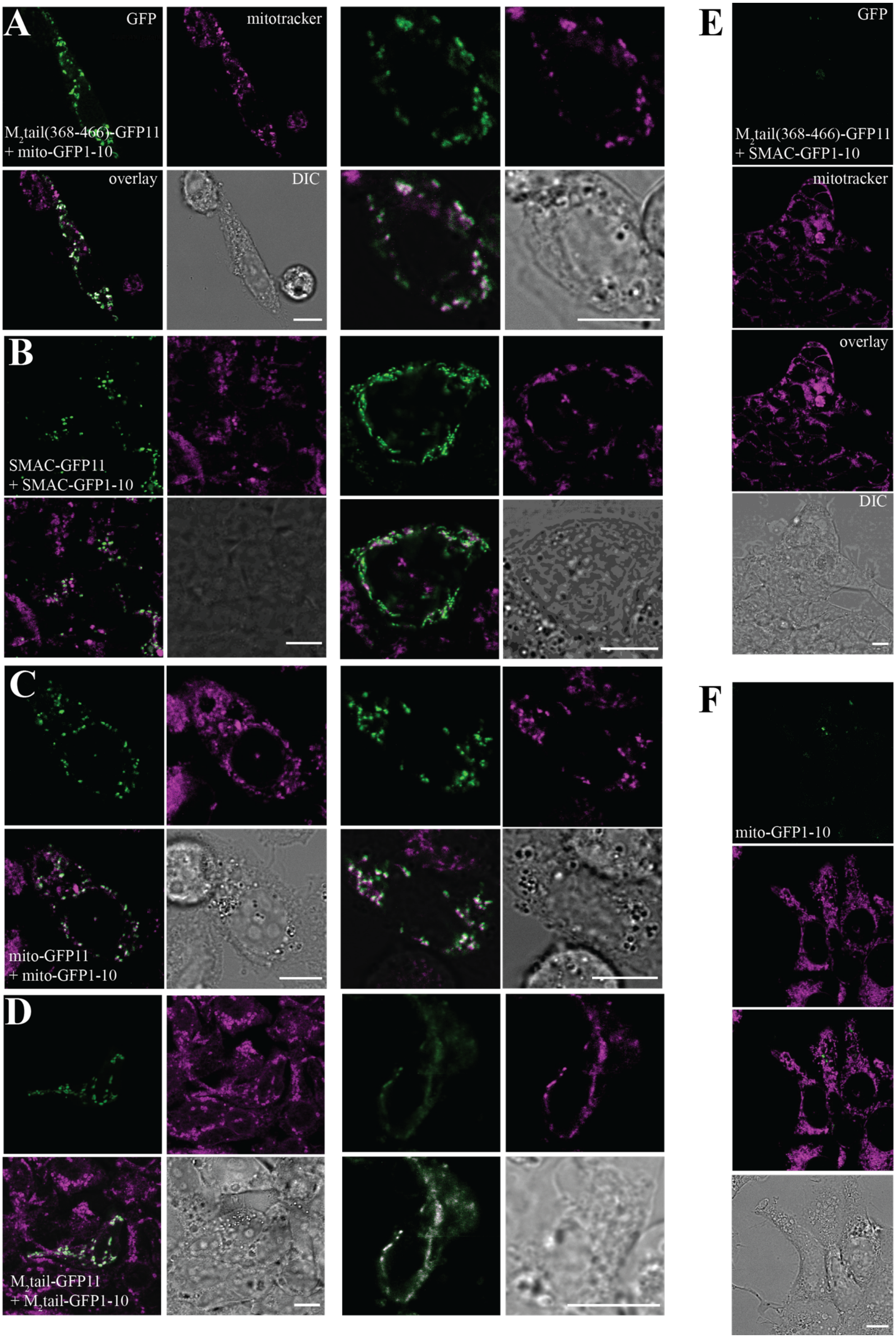
A
GFP
mitotracker
$M_2$tail(368-466)-GFP11
+ mito-GFP1-10
overlay
DIC
B
SMAC-GFP11
+ SMAC-GFP1-10
C
mito-GFP11
+ mito-GFP1-10
D
$M_2$tail-GFP11
+ $M_2$tail-GFP1-10
E
GFP
$M_2$tail(368-466)-GFP11
+ SMAC-GFP1-10
mitotracker
overlay
DIC
F
mito-GFP1-10

**Fig. S8. Cellular localization of split GFP constructs in HEK293 cells.**

**A,** Confocal images displaying the mitochondrial localization of $M_2$tail(368-466)-GFP11 and mito-GFP1-10. **B,** co-expression of SMAC-GFP11 and SMAC-GFP1-10 (positive control) and corresponding Mitotracker deep red image. **C,** mito-GFP11 and mito-GFP1-10 (positive control). **D,** $M_2$tail-GFP11 + $M_2$tail-GFP1-10, with corresponding DIC image. Two representative experiments are displayed in panels A-D out of n=5 transfections for each condition. **E,** $M_2$tail(368-466)-GFP11 and SMAC-GFP1-10. **F,** mito-GFP1-10 alone (negative control). One representative experiment is shown in panels E-F out of n04 transfections for each condition. EGFP (green) was excited at 488 nm and fluorescence collected between 500-600 nm. Mitotracker deep red (magenta) was excited at 633 nm and fluorescence collected between 650-750 nm. Scale bars are 10 µm.

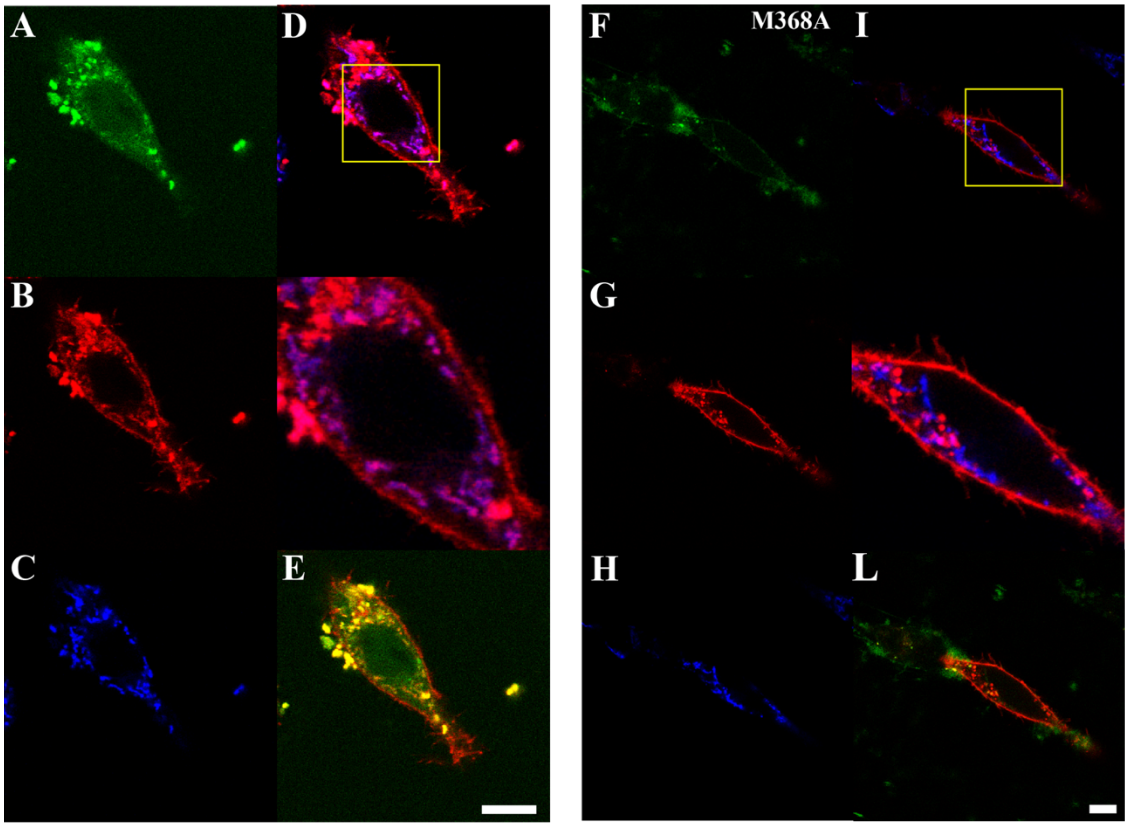


**Fig. S9. Mitochondrial localization of the $M_2$tail(368-466) fragment upon IRES production.**

**A-E**, Mitochondrial localization of the IRES mediated C-terminal fragment is enhanced upon cellular stress. HEK293 cells transfected with the super construct $M_2$-mRuby2-STOP-$M_2$-i3-tail-EGFP display increased localization of the EGFP labelled portion to the mitochondria, upon serum starvation for 2 hours and incubation of the cells in HBSS buffer, in presence of Mitotracker: A, is IRES-driven $M_2$tail(368-466)-EGFP. **B**, Cap dependent $M_2$-mRuby2 + IRES-driven $M_2$tail(368-466)-mRuby2, both derived from the $M_2$-mRuby2 gene. **C**, Mitotracker Deep Red **D**, Overlay of Mitotracker Deep Red and mRuby channel (colocalization in magenta). Zoom-in of yellow square immediately below the panel. **E**, Overlay between GFP and mRuby2 (colocalization in yellow). **F-L**, Localization of the mutant construct $M_2$(M368A)-mRuby2-STOP-$M_2$-i3-tail-EGFP in HEK293 cells after 2 hours incubation in HBSS buffer. The same legend as in **A-E** applies. Bottom: Schematic structure of the mega constructs $M_2$(M368A)-mRuby2-STOP-$M_2$-i3-tail-EGFP. This construct is similar to the $M_2$-mRuby2-STOP-$M_2$-i3-tail-EGFP construct but the methionine 368 in the $M_2$ sequence has been replaced with alanine. The asterisks represent the stop codons at the end of mRuby2 and EGFP. EGFP was excited at 488 nm and fluorescence collected between 500-600 nm; mRuby2 was excited at 561 nm and fluorescence collected between 580-620 nm. Mitotracker deep red was excited at 633 nm and fluorescence collected between 650-750 nm. Scale bars are 10 µm.

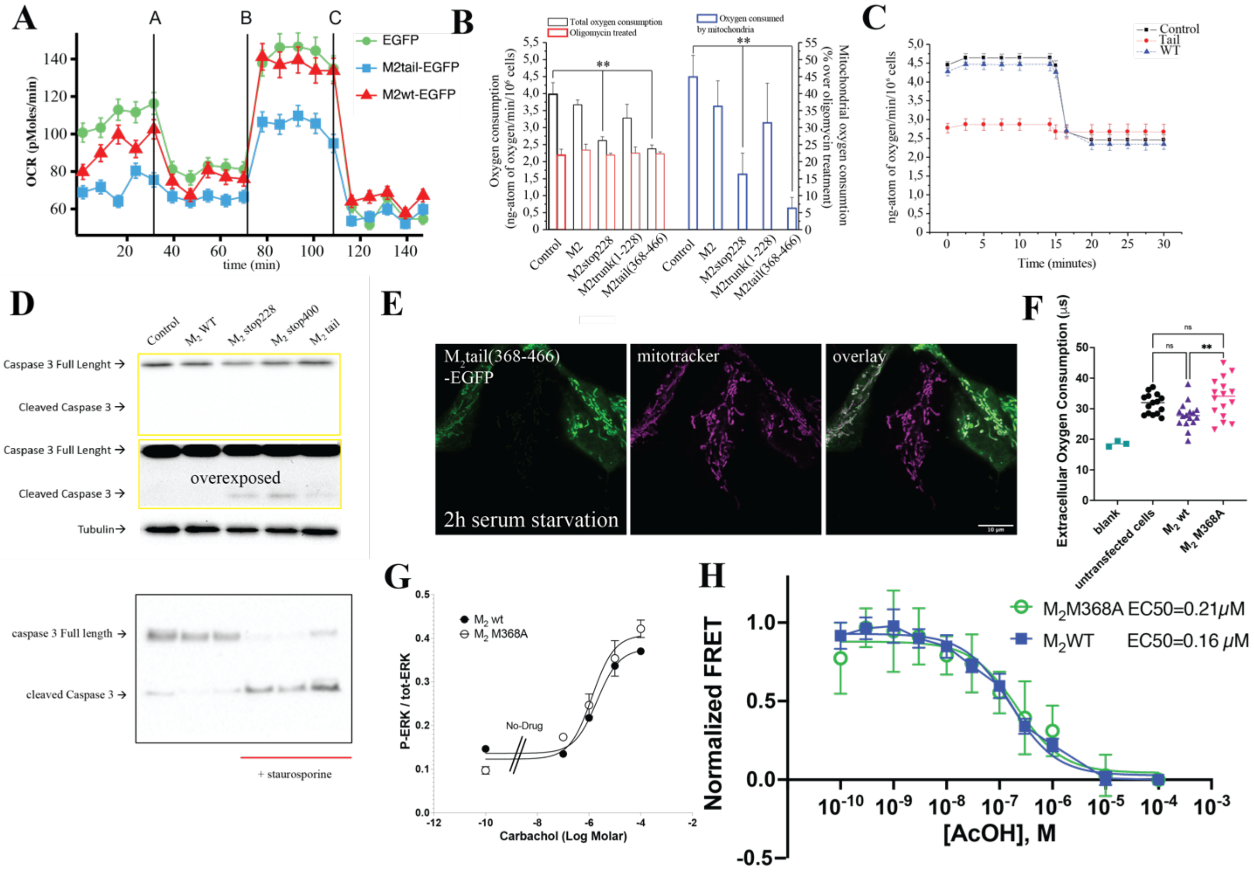


**Fig. S10. Supplemental data on the effect of the $M_2$tail(368-466) fragment on mitochondrial function and cell metabolism.**

**A,** Representative experiment of Seahorse assay showing the time course of oxygen consumption inhibition by $M_2$tail(368-466)-EGFP and wild type $M_2$-EGFP, compared to the same experiments performed in control cells transfected with EGFP. The rate of ATP-linked respiration measured after oligomycin administration is decreased in $M_2$tail and slightly decreased in $M_2$ cells as well as compared to the controls. FCCP, an ionophore, was added to each cell type to investigate the cellular bioenergetic reserves that seem to be slightly reduced only in the $M_2$ tail cells. **B,** Oxygen consumption measured by a Clark type electrode-based-polarographic method at a constant temperature of 37 °C. After 15 minutes of basal respiration, the ATP synthase inhibitor oligomycin was added at a concentration of 17 nM. Oxygen consumption in the absence and presence of oligomycin is represented by black and red bars, respectively. The blue bars represent the percentage of mitochondrial oxygen consumption (total minus oligomycin treated) in cells transfected with each construct. Significance values reported in the graphs were determined by a one-tailed Student t-test, p-values: ** $0.001<p<0.01$. **C,** is a representative experiment showing the time course of inhibition of oxygen consumption by $M_2$tail(368-466), compared to the same experiments performed in control cells and cells transfected with the $M_2$ wild type construct, upon addition of oligomycin. Results are given as ng-atom of oxygen/min/106 cells. **D,** Western blot of caspase-3 cleavage in COS-7 cells transfected with the indicated $M_2$ receptor mutants 48h after transfection (top of panel, normal contrast; bottom of panel, increased contrast). Below, as a positive cleavage control, the blot for caspase-3 with and without Staurosporine incubation. **E,** Confocal sequential images of $M_2$tail(368-466)-EGFP (green), Mitotracker (magenta) and overlay upon 2h of serum starvation. Scale Bars are 10 µm. **F,** Extracellular oxygen consumption assay of HEK293 cells transfected with the indicated $M_2$ receptor mutants using a fluorogenic dye quenched by Oxygen. Blank indicates the dye lifetime measured in wells containing the dye, but no cells. **G,** Normalized dose response to Carbachol stimulation of HeLa cells, comparing $M_2$ wild type to the $M_2$ M368A mutant. **H,** Normalized dose response curves based on FRET Gi biosensor[10], reflecting Galphai3 activation by two $M_2$ receptor mutants ($M_2$ wild type n=7 transfections, $M_2$ M368A n=3 transfections) stimulated by acethylcholine (AcOH) in HEK293 cells. Source data for panels A,B, C, F, G and H can be found in S1 Data.

**Table S1.** Ligand binding and functional properties of $M_2$ receptor mutants with single and double stop codons. Number of [$^3$H]NMS binding sites for the indicated mutants. Untransfected COS-7 cells did not show any [$^3$H]NMS specific binding. N.B. = no specific [$^3$H]NMS binding.

| Receptor | Bmax (fmol/mg) | Receptor | Bmax (fmol/mg) |
|---|---|---|---|
| $M_2$stop228 | 45.3 ± 2.4 | $M_2$stop228/stop368 | N.B. |
| $M_2$stop196 | N.B. | $M_2$stop228/stop368 + $M_2$tail(281-466) | 370± 20 |
| $M_2$stop400 | N.B. | $M_2$stop228/stop400 | N.B. |
| $M_2$stop196+ $M_2$trunk(1-283) | 47 ± 3 | $M_2$stop228/stop400+ $M_2$tail(281-466) | 380 ± 20 |
| $M_2$stop400+ $M_2$tail(281-466) | 400 ± 30 | $M_2$stop196/stop400+ $M_2$trunk(1-283) | N.B. |
| $M_2$stop196/stop400 | N.B. | $M_2$stop196/stop400+ $M_2$tail(281-466) | N.B. |
| $M_2$stop228/stop248 | 49 ± 5 | $M_2$stop228/fr.sh. | 47 ± 3 |
| $M_2$stop228/stop296 | 38 ± 3 | $M_2$stop228/Hairpin | 37 ± 2 |

**Table S2.** Radioligand binding and activation properties of $M_3$ receptor mutants. Binding characteristics and stimulation of phosphatidylinositol accumulation by $M_3$, $M_3$stop273 and the co-transfected $M_3$trunk(1-272) and $M_3$tail(M-388-586). Untransfected COS-7 cells did not show any [$^3$H]NMS specific binding. N.B. = no specific [$^3$H]NMS binding.

| Binding data | | | | Phosphatidylinositol accumulation assay | |
|---|---|---|---|---|---|
| Receptor | Bmax (fmol/mg) | [$^3$H]NMS KD (pM) | Carbachol IC50 (μM) | Carbachol EC50 (μM) | Maximum increase in IP1 level above baseline (%) |
| $M_3$ | 1120 ± 71 | 29 ± 2 | 59 ± 4 | 4.2 ± 0.5 | 183 ± 15 |
| $M_3$trunk(1-272) + $M_3$tail(388-589) | 114 ± 17 | 23 ± 2 | 48 ± 5 | 1.4 ± 0.1 | 126 ± 12 |
| $M_3$stop 273 | 52.3 ± 0.5 | 31.3 ± 2.8 | 56 ± 2 | 1.1 ± 0.1 | 89 ± 13 |

**Table S3**. Ligand binding of $M_3$ receptor mutants with single and double stop codons. Number of [$^3$H]NMS binding sites in COS-7 cells transiently transfected with the indicated $M_3$ receptor mutants with single and double stop codons.

| Receptor | Bmax (fmol/mg) | Receptor | Bmax (fmol/mg) |
|---|---|---|---|
| $M_3$stop273 | 54 ± 1 | $M_3$stop273/stop503 | N.B. |
| $M_3$stop240 | N.B. | $M_3$stop273/stop503 + $M_3$tail(388-589) | 120± 15 |
| $M_3$stop503 | N.B. | $M_3$stop240/stop503+ $M_3$trunk(1-272) | N.B. |
| $M_3$stop240+ $M_3$trunk(1-272) | 44 ± 3 | $M_3$stop240/stop503+ $M_3$tail(388-589) | N.B. |
| $M_3$stop503+ $M_3$tail(388-589) | 135 ± 12 | $M_3$stop273/fr.sh. | 120 ± 4 |
| $M_3$stop240/stop503 | N.B. | $M_3$stop273/Hairpin | 30 ± 2 |
